# MSFragger-Labile: A Flexible Method to Improve Labile PTM Analysis in Proteomics

**DOI:** 10.1101/2022.10.12.511963

**Authors:** Daniel A. Polasky, Daniel J. Geiszler, Fengchao Yu, Kai Li, Guo Ci Teo, Alexey I. Nesvizhskii

## Abstract

Post-translational modifications of proteins play essential roles in defining and regulating the functions of the proteins they decorate, making identification of these modifications critical to understanding biology and disease. Methods for enriching and analyzing a wide variety of biological and chemical modifications of proteins have been developed using mass spectrometry (MS)-based proteomics, largely relying on traditional database search methods to annotate resulting mass spectra of modified peptides. These database search methods treat modifications as static attachments of a mass to particular position in the peptide sequence, but many modifications undergo fragmentation in tandem MS experiments alongside, or instead of, the peptide backbone. While this fragmentation can confound traditional search methods, it also offers unique opportunities for improved searches that incorporate modification-specific fragment ions. Here, we present a new Labile Mode in the MSFragger search engine that can tailor modification-centric searches to the fragmentation observed. We show that labile mode can dramatically improve spectrum annotation rates of phosphopeptides, RNA-crosslinked peptides, and ADP-ribosylated peptides. Each of these modifications presents distinct fragmentation characteristics, showcasing the flexibility of MSFragger labile mode to improve search for a wide variety of biological and chemical modifications.

## Introduction

Post-translational modifications (PTMs) of proteins play essential roles in defining and regulating protein functions(1). Liquid chromatography-tandem mass spectrometry (LC-MS/MS) methods have become the preferred method for large-scale identification of proteins from biological samples, becoming known as mass spectrometry (MS)-based proteomics(2). The ability of MS to detect and characterize post-translational and chemical modifications of proteins at whole-proteome scales is currently unmatched and provides crucial insights into the function and regulation of proteins in biological systems. Analysis of PTMs is typically accomplished by broadly similar MS methods as those used for unmodified peptide analysis in proteomics, often with an added enrichment step to concentrate modified peptides of interest prior to LC-MS/MS analysis. Database search methods to annotate the resulting tandem mass spectra are also similar to those used for unmodified peptides, as indeed, chemical artifacts and common modifications are nearly always included in such standard proteomics searches. The predominant search strategy for modified peptides assumes that modifications will remain intact on the peptide, simply shifting the mass of the amino acid to which they are connected by a fixed value(3). Fragment ions matching the combined amino acid and modification masses are used to detect modified peptides and localize the modification within the peptide. However, many modifications experience their own fragmentation during tandem MS, causing a mismatch between the expected and observed fragment ions. Some peptides can still be identified, by matching only ions that do not contain the modification site, but many spectra of modified peptides cannot be annotated, contributing to the “dark matter” of proteomics(4, 5).

Labile modifications, or those that fragment instead of or in addition to peptide backbone fragmentation in tandem MS, are extremely common. The most abundant PTMs, phosphorylation(6–8) and glycosylation, are labile, along with many less abundant PTMs like sulfation(9) and ADP-ribosylation(10), and many other peptide modifications of interest, including chemical modifications used for chemoproteomics methods(11), ubiquitinylated and SUMOylated peptides retaining a piece of a second peptide after digestion, RNA-crosslinked peptides(12), and many more. As many modifications of interest are labile, several approaches have been devised to identify and localize peptides bearing labile modifications. Electron-based activation methods like ETD and ECD typically(13), though not always(14), preserve modifications intact even if they are labile in collision activation methods, allowing database search methods to confidently identify modified peptides(15, 16). However, these methods come at the cost of reduced acquisition speed(13), leading many groups to continue utilizing collisional activation for PTM searches. Particularly for modifications that are only partially labile, such as phosphorylation, collisional activation remains the most common analysis method. Many traditional search engines allow specified neutral losses from a modification, such as loss of phosphoric acid from phosphorylated peptides(17–19), which allows for improved search of modified peptides in some cases but lacks the flexibility to handle more complex modifications, such as glycosylation. ProteinProspector additionally allows searching for any modification that is lost entirely during fragmentation(18). For the most part, however, PTM search tools either utilize no modification-specific fragments beyond a few common neutral losses or require bespoke search methods tailored to a particular modification, as is common for glycosylation.

Recently, an alternative approach has emerged in the “open” search method developed for searching for unknown or unexpected peptide modifications(4, 5). Open searches, and the similar mass offset or multinotch searches(20), allow searching of peptides with a “mass offset,” or a difference between the peptide sequence mass and the observed precursor mass from MS^1^. This allows peptides with unknown (open search) or known (offset search) modifications to be identified. Crucially for searching labile modifications, the peptide sequence is often initially searched without modifications in these methods, as a consequence how fragment-ion indexing approaches are implemented to improve the speed of such searches. As this reduces the sensitivity of the search for nonlabile modifications, a “localization-aware” open search method was recently implemented in MSFragger to allow searching for fragment ions retaining intact modification(21). However, the default open/offset search is a natural fit for identifying labile modifications that are lost completely from the peptide during MS^2^, as searching for peptide fragment ions without the modification is the optimal method in these cases. The MSFragger Glyco search method took advantage of this for searching glycopeptides, dramatically boosting sensitivity by searching for just peptide backbone fragment ions without glycosylation(22). Many modifications are not lost so cleanly, however, either leaving behind a piece of the modification, or taking a piece of the peptide with them when leaving, such as the net loss of water from phospho-Ser and Thr residues during neutral loss of phosphoric acid. Modification loss can also produce signature, or diagnostic, fragment ions in the mass spectrum that indicate the presence of a particular modification(23).

To support a wide range of labile modification searches, we have implemented several new features in MSFragger, referred to collectively as “labile mode” search. Labile mode searches allow filtering of spectra for diagnostic ions when considering a labile modification, as well as specifying peptide or fragment remainder masses resulting from modification fragmentation to improve spectrum annotation and localization. Here, we demonstrate the utility of labile mode searches to increase the spectral annotation rate of data for several labile modifications, including phosphorylation, peptide-RNA crosslinks, and ADP-ribosylation. We used our recently described diagnostic ion mining module in PTM-Shepherd to determine the fragment ions to use in labile search, allowing optimal settings to be chosen even for modifications without well characterized fragmentation pathways(23). Labile search has been incorporated into MSFragger starting with version 3.5 and FragPipe version 18.0, and workflow templates for the labile phosphorylation and ADP-ribosylation searches described here are available from the workflows menu in FragPipe.

## Experimental Procedures

### Datasets

“RNA crosslinking” data was downloaded from PXD023401, from the report of Bae *et al*(12). Briefly, crosslinked RNA-peptide complexes were prepared by photoactivable ribonucleoside labeling and UV crosslinking and analyzed by LC-MS/MS on an Orbitrap Fusion Lumos instrument using HCD activation at 30% normalized collision energy (NCE). Data corresponding to the 4-thiouridine (4SU) crosslinked RNA-peptide complexes were analyzed here. “CPTAC” phosphoproteomics data from the Clear Cell Renal Cell Carcinoma (CCRCC) cohort was downloaded from the Clinical Proteomic Tumor Analysis Consortium (CPTAC) data portal. TMT-labeled phosphopeptides were enriched from tumor and adjacent normal tissue samples, fractionated, and analyzed on an Orbitrap Fusion Lumos mass spectrometer using HCD activation at 37 NCE(24). “Multi-energy” phosphoproteomics data was downloaded from PXD004415 and refers to the report of Tran *et al*., who analyzed unlabeled, enriched phosphopeptides from Jurkat T cells via HCD at two collision energies, 25 and 35 NCE, on an Orbitrap Q Exactive mass spectrometer(25). “AIETD” ADP-Ribosylation data was downloaded from PXD017417, in which ADP-Ribosylated peptides were enriched using Af1521 macrodomain affinity from HeLa cells treated with H2O2 and analyzed on an Orbitrap Fusion Lumos instrument modified with a CO2 laser for AIETD experiments(26). “HCD” ADP-Ribosylation data was downloaded from PXD004245, where ADP-Ribosylated peptides from H2O2-treated HeLa cells and mouse liver tissues were enriched using Af1521 macrodomain and analyzed by HCD activation on an Orbitrap Q Exactive instrument at 28 NCE(27).

### General Search Settings and Validation

For all searches, raw files were converted to mzML format and centroided using MSConvert version 3.0.22068 from ProteoWizard(28, 29). All searches were performed using MSFragger version 3.5, Philosopher version 4.2.2, and FragPipe version 18.0. Several options are available for validation of labile search results. In all searches reported here, labile modifications from mass offset searches were written to the MSFragger output in the same manner as variable modifications from conventional (nonlabile) MSFragger searches using the “Report mass shift as a variable mod” option, and downstream validation tools were used as in conventional searches. For open searches and searches with many modifications, labile modifications can instead be left as mass offsets (reported as “delta masses” by MSFragger) and the extended mass model in PeptideProphet(30) can be used to model probabilities for each mass offset, as is done in MSFragger Glyco searches(22). For result scoring, Percolator(31) or PeptideProphet can be used in FragPipe. Percolator is now supported for open searches from MSFragger following implementation of a delta mass score in FragPipe, however, PeptideProphet is still recommended as the extended mass model provides more rigorous treatment of delta masses. For mass offset labile searches, either Percolator or PeptideProphet can be used. For localization of modifications, PTMProphet(32) is available in FragPipe in addition to the fragment remainder-based localization of MSFragger and can be used instead of or in addition to MSFragger localization. All searches performed here used ProteinProphet(33) for protein inference and performed FDR filtering to 1% PSM, ion, peptide, and protein levels in Philosopher using the sequential method(34).

### RNA-XL search

4SU data was searched against a reviewed human proteome with decoys and common contaminants added in Philosopher, downloaded 2022/06/13 with 20,420 total entries (including contaminants). Fully enzymatic MSFragger search was performed using strict-trypsin settings *(i.e.,* allowing cleavage before Pro) and allowing 2 missed cleavages, precursor and fragment tolerances of 20 and 10 ppm respectively, mass calibration, fixed Cys carbamidomethylation, and variable modifications of Met oxidation and Protein N-terminal acetylation. 4SU crosslinking was searched as both the intact nucleoside (+226.0594 Da) and the base only (+94.0168 Da), as frequent in-source fragmentation of nucleoside was noted by Bae *et al.* and which we confirmed using the PTM-Shepherd diagnostic mining module(23). For labile search, +94 and +226 were searched as mass offsets on any amino acid, with labile mode active, no diagnostic or peptide remainder ions, and a fragment remainder ion of +94. For the equivalent nonlabile search, variable modifications of +94 and +226 were specified on all amino acids (max of 1 per peptide). PeptideProphet was used for validation with default closed search settings.

### CPTAC phosphoproteomics search

The CCRCC cohort phosphoproteomics data were searched against the same human database as used in the RNA-XL search. MSFragger search was performed with fully enzymatic strict-trypsin search allowing 2 missed cleavages, precursor and fragment tolerances of 20 and 10 ppm respectively, mass calibration, fixed modifications of Cys carbamidomethylation and TMT-labeling of Lys and peptide N-termini, and variable modifications of Met oxidation, Protein N-terminal acetylation, deamidation (Asn, Gln), and peptide N-terminal pyro-Glu (Gln, Cys). Up to 3 phosphorylation events were allowed on a peptide, specified as variable modifications (nonlabile search) and/or mass offsets (labile search), with the number of nonlabile and labile phosphorylations always adding up to 3. For labile searches, a fragment remainder ion of −18.01056 was specified, corresponding to neutral loss of phosphoric acid, but no diagnostic or peptide remainder ions. PeptideProphet was used for validation with default closed search settings, combining all fractions of each TMT-plex for modeling together.

### Multi-Energy phosphoproteomics search

All data from Tran *et al.* at both HCD settings (25 and 35) were searched against the same database as CPTAC search, with identical MSFragger and validation settings except for no TMT labeling.

### ADP-Ribosylation searches

Searches of HeLa data used the same human database as the RNA-XL and phosphoproteomics searches. Searches of Mouse tissue data used a reviewed mouse proteome with decoys and common contaminants added in Philosopher, downloaded 2022/06/27 with 17,230 total entries. MSFragger searches were fully enzymatic, using Trypsin settings with up to 5 missed cleavages, precursor and fragment tolerances of 20 and 10 ppm respectively, mass calibration, fixed modification of Cys carbamidomethylation, variable modifications of oxidation (Met, Trp), protein N-terminal acetylation, and peptide n-terminal pyro-Glu (Gln, Cys). For labile searches, ADP-Ribose (+541.06111 Da) was specified as a mass offset on Ser (HeLa) or Ser, Arg, and Lys (mouse) with diagnostic ions (136.06232, 250.09401, 348.07036, 428.03669 Da for all searches and additionally 584.09018 Da for mouse tissue only) and peptide remainder ions (114.03169, 193.99802, 291.97492, and 406.00661 Da). A fragment remainder mass of −42.0205 Da was specified for mouse tissue only (corresponding to partial loss of the Arg side chain). Nonlabile searches specified ADP-Ribose as a variable modification of Ser (HeLa) or Ser, Arg, and Lys (mouse). HCD searches used peptide *b* and *y* ions only; AIETD searches used *b, y, c,* and *z* ions. Percolator was used for scoring with default parameters and mass offset score from FragPipe, after testing indicated improved performance over PeptideProphet for this dataset.

### Experimental Design and Statistical Rationale

No experiments were performed for this study; all results are derived from reanalysis of previously published datasets (for details, see “Datasets”).

## Results and Discussion

Many peptide modifications undergo fragmentation in tandem mass spectrometry experiments, changing the expected mass of peptide backbone fragment ions and resulting in poor performance for traditional search methods. The MSFragger labile search mode provides a flexible and general approach to capture the ions resulting from modification fragmentation, allowing spectra to be filtered for diagnostic ions and searched for partial or complete fragmentation of a modification with peptide and fragment remainder ions. To demonstrate the utility of the labile search method, we re-analyzed published data corresponding to three PTMs, each with a distinct fragmentation pattern that is supported by the labile search mode.

An overview of the ion types and methods included in labile mode is shown in Figure 1. MSFragger labile mode search is an extension of the MSFragger search engine’s(5) open and mass offset search modes for labile modifications, and consists of three main features corresponding to the three types of fragment ions generated by fragmentation of labile modifications (Fig. 1A). First, “diagnostic ions,” or peaks corresponding to a portion of the modification observed on its own after dissociation from a modified peptide, can be specified along with a minimum intensity threshold (Fig. 1A, left). Diagnostic ions indicate spectra containing a modified peptide and are used as a filter for labile mode searches, in which only spectra where the sum of intensities of the diagnostic ions exceeds a user-specified threshold are searched for the labile modification(s) (*i.e*., mass offsets). All other spectra not containing sufficient diagnostic ion signal are searched only for peptides with no mass offset (*i.e*., unmodified peptides). Second, “peptide remainder ions,” or the intact peptide bearing a partially fragmented modification, can be specified (Fig. 1A, right). Peptide remainder ions are common for larger modifications containing multiple labile bonds, such as the ADP-Ribosylation depicted in Fig. 1. In labile searches, these ions are added to the peptide score, providing a boost for modified peptides that produce them. Finally, “fragment remainder ions” (Fig. 1A, center) refer to peptide backbone fragment ions that retain a part of the modification. Fragment remainder ions are added to the peptide score like peptide remainder ions, boosting sensitivity for modified peptides. Unlike peptide remainder ions, fragment remainder ions are also used to localize the modification within the peptide sequence, since only fragments that include the modification site can generate a modified ion. Negative fragment remainders can also be specified, referring to loss of part of the amino acid side chain at the modification site during loss of modification, such as loss of phosphoric acid from phospho-Ser or Thr residues.

**Figure 1.**
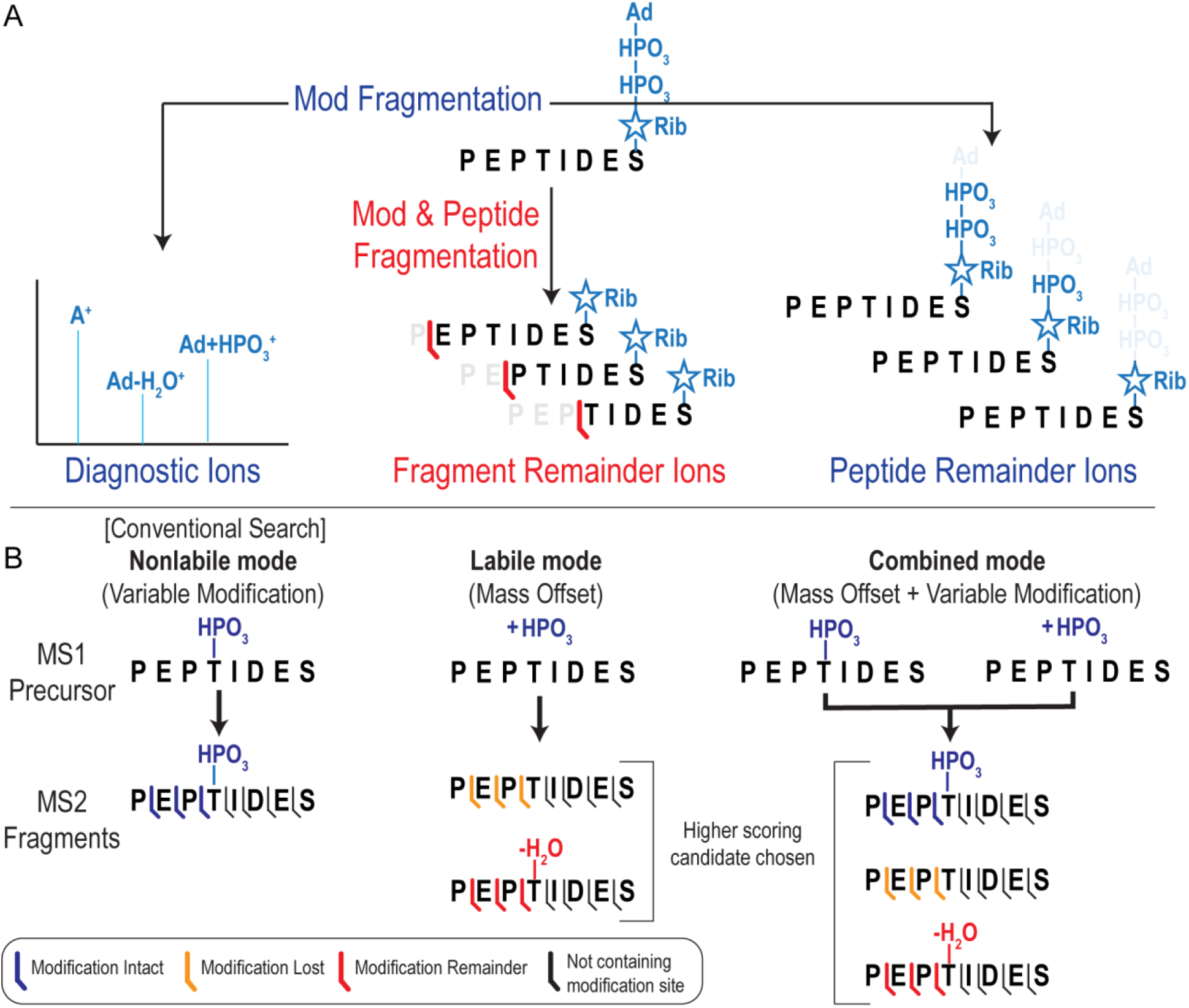
MSFragger labile mode workflows. A) Modification fragment types available for labile mode searches, including diagnostic ions (partial modification only), peptide remainder ions (intact peptide with partial modification), and fragment remainder ions (partial peptide with partial modification). B) MSFragger search modes. In nonlabile mode, modifications are specified as variable modifications (as in conventional MSFragger searches). In labile mode, modifications are specified as mass offsets and fragments lacking the modification (orange) are searched. If fragment remainders (red) are specified, candidates with modification loss ions and fragment remainder ions are compared, with the higher scoring candidate chosen. In combined mode, a modification is specified as both a variable modification and mass offset, producing labile and nonlabile candidates. As in the labile mode, the highest scoring candidate is chosen as the search result.

MSFragger searches can now be performed entirely in the conventional (nonlabile) mode, entirely in the new labile mode, or in a combination mode (Fig. 1B). In the combination mode, modifications that incompletely dissociate can be specified as both a variable modification (nonlabile) and as a mass offset (labile). Spectra will then be searched against a theoretical peptide bearing the intact modification (from the variable modification search) and against a theoretical peptide that has lost or fragmented the modification (from the labile mass offset search), with the highest scoring result being chosen as the annotation for each spectrum. The combined search will also consider peptides bearing both a variable modification and a mass offset, for a total of two modifications on the peptide. Finally, labile search results can be examined using the integrated FragPipe visualization tool FP-PDV(35). Support for viewing custom remainder fragment ions of peptides has been added, allowing examination and comparison of different fragmentation pathways directly within the graphical interface.

A recent report of a method for photoactivatable ribonucleosides for finding sites of RNA-protein crosslinks (RNA-XL) makes an excellent example of the importance of labile mode search for highly labile modifications. In this method, a single ribonucleoside is crosslinked to peptides prior to tandem MS analysis. The ribonucleoside is highly labile, exhibiting both neutral loss of ribose at low collision energy, leaving the base behind, and complete loss of the entire ribonucleoside at moderate to high collision energies(12). As a result, typical nonlabile search methods struggle to identify RNA-crosslinked peptides because fragment ions including the modification site, expected to contain the full ribonucleoside, are not found (Fig. 2A, top spectrum). With MSFragger’s labile search, fragment remainder ions (retaining the base) provide a confident identification by matching many more ions in the spectrum and allowing for localization of the modification site (Fig. 2B, bottom spectrum). We compared a nonlabile search in MSFragger (the ribonucleoside specified as a variable modification) to a two labile mode searches, one assuming complete loss of the ribonucleoside (labile mode with no fragment remainder ions) and one including a +94 Da fragment remainder corresponding to retention of the base. The labile mode searches dramatically outperformed the nonlabile search, annotating 48% and 109% more RNA-XL spectra for the no-remainder and with-remainder searches, respectively (Fig. 2B). Workaround methods have been used to search modification losses in conventional search engines, such as the method of spectrum duplication and manual precursor mass adjustment used by Bae *et al*. to search these data(12). In contrast, MSFragger labile mode provides greater capabilities, such as the remainder fragment ions that dramatically boost the number of crosslinked spectra annotated, as well as the flexibility to combine multiple such features in a fully automated fashion. Ultimately, this improved spectrum annotation translated to confident annotation of 60% more unique RNA-crosslinked peptides from 25% more proteins compared to a conventional (nonlabile) search performed with MSFragger (Fig. 2C). For highly labile modifications like these RNA-crosslinked peptides, and as previously demonstrated for glycopeptides(22), labile search thus greatly improves our capability to confidently identify modified peptides due to the near complete absence of ions bearing the intact modification that a nonlabile search requires. Fragment remainder ions are particularly advantageous because they enable localization of the modification within the peptide, analogously to how intact modifications are localized in nonlabile searches.

**Figure 2.**
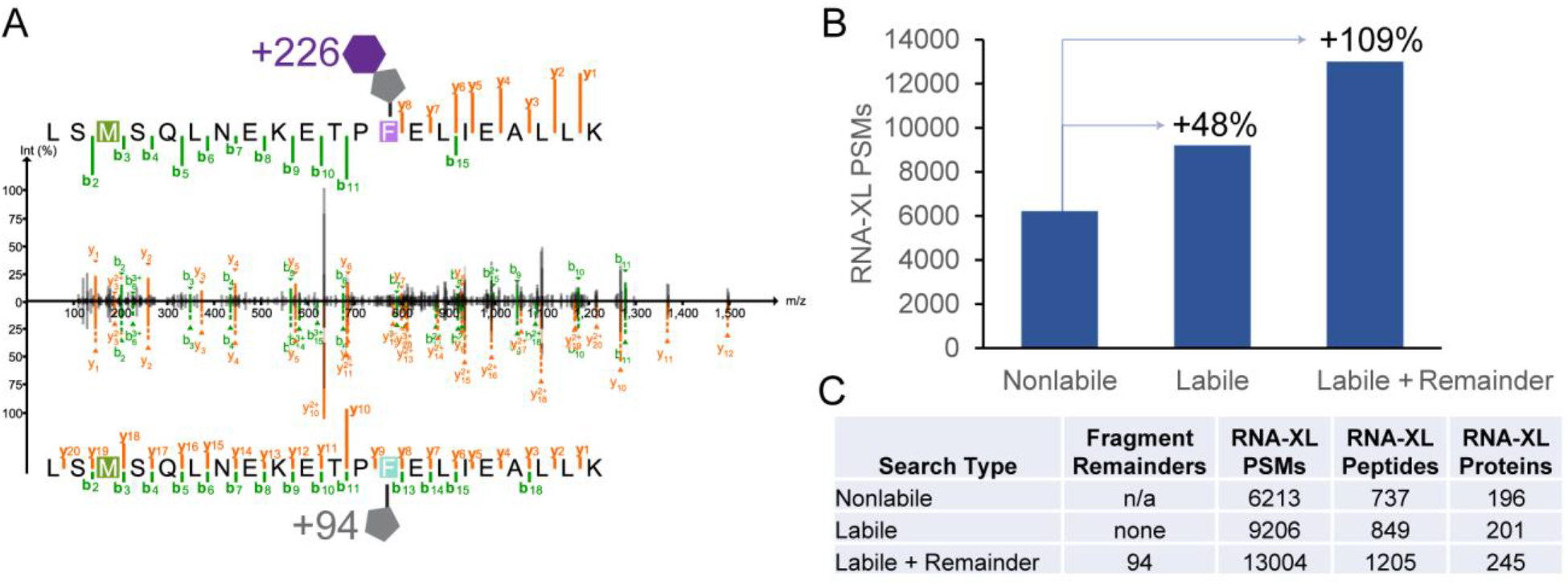
RNA-crosslinking labile search results. A) Comparison of nonlabile search (top) where fragment ions retain intact modification (mass of 226 Da) and labile search (bottom) where fragment ions retain a partial modification (mass of 94 Da). Fragment ions containing the modification site are only matched using the labile mode search. B) RNA-crosslinked PSMs found in nonlabile, labile (without remainder fragment), and labile with remainder fragment searches. C) Table of RNA-crosslinked PSMs, peptides, and proteins for the same search methods as in (B).

Not all modifications are as labile as these RNA-crosslinks, but labile search can still provide substantial benefits for intermediately labile modifications through combined searches. Phosphorylation is one of the most widely studied PTMs, with phosphoproteomics comprising an entire subfield of proteomics. It is well known that phosphorylation, particularly of Ser and Thr, can lose phosphoric acid in tandem MS, especially under conditions of low proton mobility and at high collision energies(7, 8). Despite this, many phosphopeptide searches are performed with conventional (nonlabile) methods, though some search engines support specifying the neutral loss of phosphoric acid. As peptides are often observed at multiple charge states and can vary widely in their ability to sequester protons, a mix of phosphate retention and neutral loss is typically observed, with the proportion of peptides in each category depending on the collision energy, charge state, and other factors affecting proton mobility. This mix of modification retention and loss is common to many intermediately labile modifications and presents a challenge to search engines, as there will be some peptides for which the observed fragmentation does not match the fragment ions expected by nonlabile searches and some not well matched to labile searches. MSFragger labile search provides a flexible solution to this problem, allowing for competition between the nonlabile and labile alternatives to obtain the best scoring match for each spectrum. This is accomplished by specifying both a variable modification (nonlabile) and a mass offset (labile) of the same mass and amino acid specificity. If the peptide in a given spectrum largely retains the modification, the nonlabile variable modification version of the peptide will obtain a better score, whereas if the peptide largely loses the modification, the labile mass offset version will be chosen instead, allowing both possibilities to be considered in a single search (Fig. 1B, combined mode). For phosphopeptide searches, peptides that contain multiple phosphates can be searched by setting more than one variable modification and/or a mass offset corresponding to the mass of multiple phosphate modifications. Mass offset searches are restricted to considering a single modification site for nonlabile modifications, but multiple labile modifications can be successfully identified (without localization) thanks to dissociation of the modifications. In the phospho searches described here, we allowed a maximum of 3 phosphates per peptide and tested each combination of variable modifications and mass offsets yielding a maximum of 3 modifications to evaluate the performance of labile, nonlabile, and combination searches without interference from searching different numbers of phosphates.

We demonstrate the value of this flexible labile search method for phosphoproteomics data from two sources, one label-free with fragmentation compared at low and high collision energies, and one TMT-labeled and fragmented at high energy. The label-free phosphoproteomics data of Tran *et al.* shows the dependence of phosphate loss on collision energy(25). At an NCE of 25, the labile and nonlabile searches have very similar performance (Fig. 3A). Combined labile/nonlabile search offers a slight improvement in performance, presumably as some peptides exhibit phosphate loss while others do not, but the difference is minimal (< 4%) compared to either fully labile or nonlabile search. At the higher NCE setting of 35, however, the fully labile search annotates over 20% more spectra than the fully nonlabile search, as the majority of phosphates are dissociated at this high energy (Fig. 3A). TMT labeling has been shown to increase phosphate neutral loss by reducing proton mobility(8), leading us to also examine a TMT-labeled phosphoproteomics dataset from the CPTAC consortium (see Methods for details). This large dataset, comprising 23 TMT-11 plexes, was searched with the same set of nonlabile, combined, and fully labile phopsho searches. Between the high collision energy (NCE 37) and TMT labeling, it is no surprise that the fully labile search offered the best performance, annotating roughly 200,000 more phosphopeptide spectra, or an increase of 11%, compared to the nonlabile search (Fig. 3B). The combined mode searches also provided substantively improved performance compared to conventional nonlabile search, possibly as a result of the variation in proton mobility from peptides with different numbers of TMT labels. The combined and fully labile searches ultimately annotated roughly 4,000 more phosphopeptides and 300 more phosphoproteins than the nonlabile search (Fig. 3C), demonstrating the value of considering phosphate loss in search, particularly for TMT-labeled phosphopeptides fragmented at high energy.

**Figure 3.**
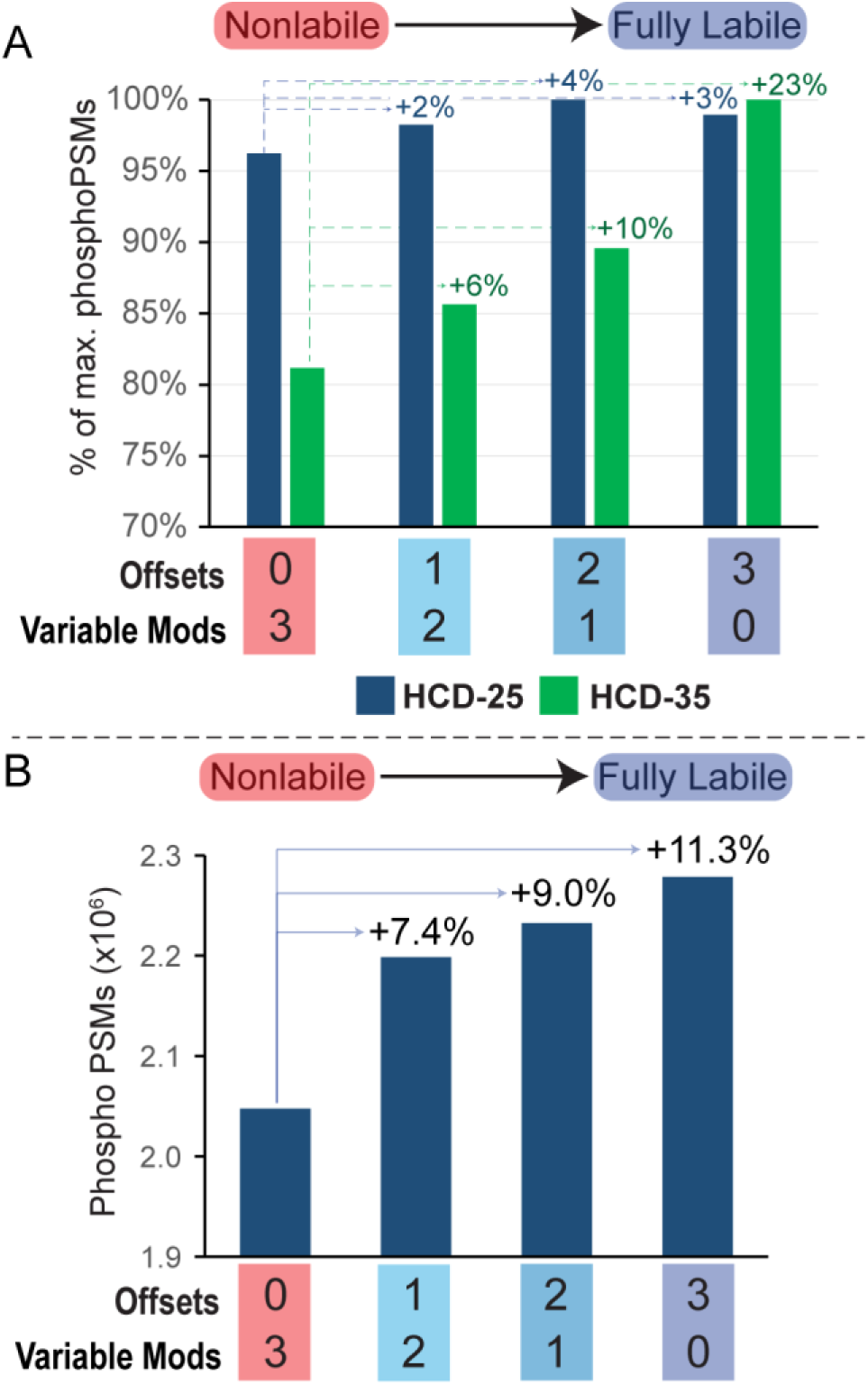
Phosphopeptide searches with labile mode. A) Data from Tran et al. searched with up to 3 phosphorylation events per peptide. Search results from left to right increase the number of labile phosphorylations vs nonlabile, from 0 (fully nonlabile) at left to 3 (fully labile) at right. PhosphoPSM counts increase substantially in labile searches of HCD-35 data (green), vs minimal difference in HCD-25 data (blue). B) PhosphoPSMs found in searches of CPTAC CCRCC cohort TMT-labeled phospho-enriched samples using the same nonlabile to labile search progression as in (A).

Finally, we re-analyzed several datasets containing ADP-ribosylated peptides to assess the performance of MSFragger labile search on a modification with different degrees of lability and from hybrid activation methods. Analysis of ADP-ribosylation is a growing field of study, and it has been established that ADP-ribose loss from Ser residues is extremely common(10, 36), leading to many current analyses employing hybrid activation methods like EThcD to localize ADP-ribosylation sites(37, 38). A recent study employed AIETD and compared a range of laser powers for activation, concluding that a medium laser power setting was best for the nonlabile search employed in the study(26). As in HCD fragmentation, we observe a strong positive correlation between laser power and the performance of the combined mode search compared to fully nonlabile search, with combined mode search providing improved performance at all but the lowest two laser powers (Fig. 4A). Given that the primary fragmentation is expected to come from electron transfer, preserving modifications intact, smaller improvements in performance than in HCD analyses are expected; however, labile search methods can clearly still provide benefits in hybrid activation methods when supplemental activation is sufficiently high-energy, or modifications are sufficiently labile. We also examined HCD activation ADP-ribosylation data from HeLa cells (primarily containing Ser-linked ADP-R) and mouse liver tissue (containing a mix of Arg- and Ser-linked ADP-R). Labile search offered a greater increase in PSMs annotated for Ser-linked ADP-R in the HeLa data, as the Ser linkage is significantly more labile (Fig. 4B, C). However, Arg-linked ADP-R uniquely generates a fragment remainder ion with a mass of −42 Da, corresponding to loss of CN2H2 from the Arg side chain(10). Adding this remainder fragment resulted in a notable increase in PSMs annotated in mouse liver tissue, but not in HeLa lysate, as expected given the different proportions of Arg-linked ADP-R (Fig. 4D, center). The presence of this fragment remainder allowed for confident localization of the majority of Arg-linked ADP-R sites in the mouse liver data (Fig. 4D, blue), whereas Ser-linked sites left no fragment remainder and generally could not be localized from HCD data in either mouse liver or HeLa lysate (Fig. 4D, red). Thus, while Ser-linked ADP-R indeed requires hybrid activation for localization, Arg-linked ADP-R can be successfully localized by the faster and more sensitive HCD activation using MSFragger labile mode.

**Figure 4.**
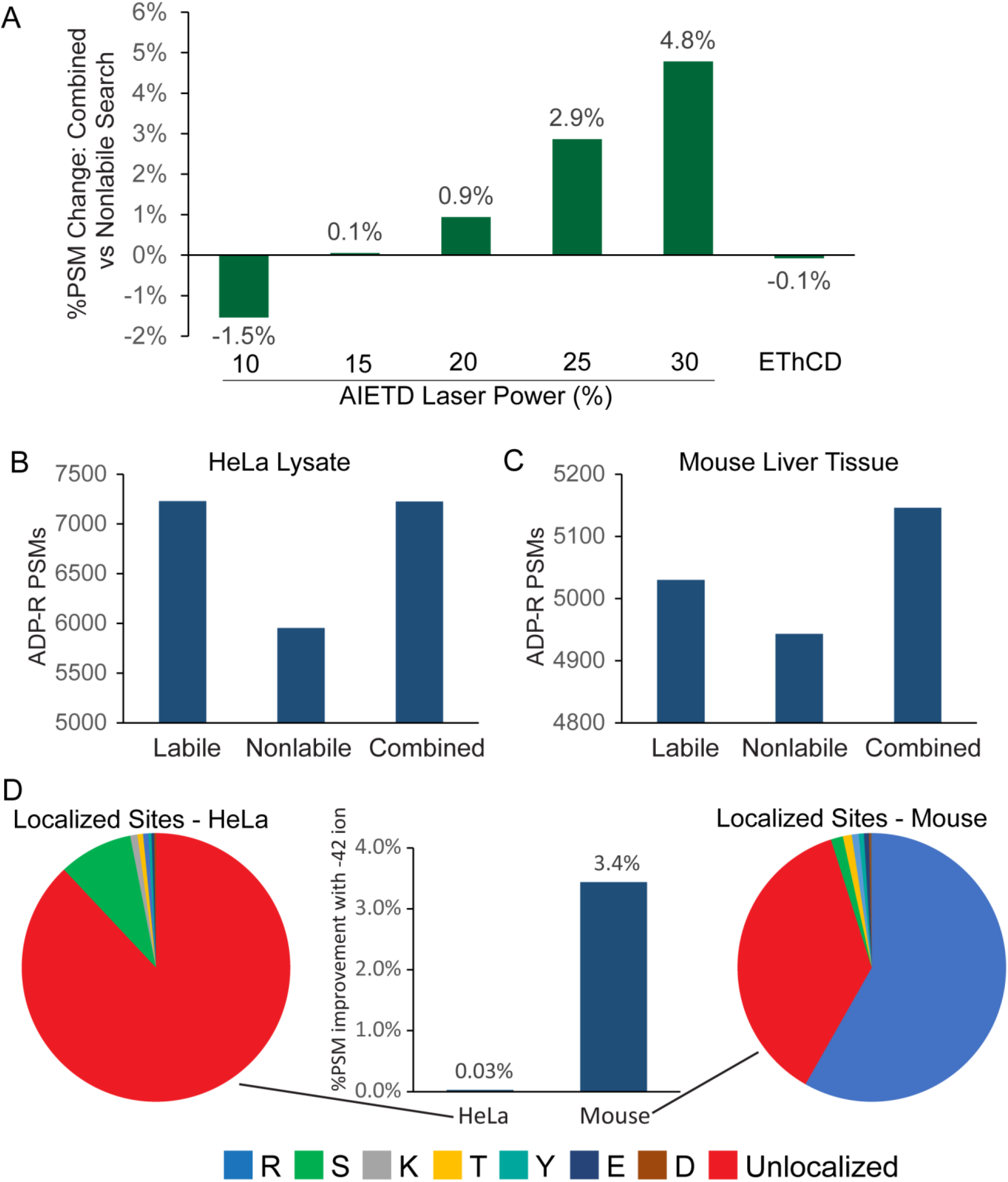
ADP-ribosylation searches with labile mode. A) Percentage increase in combined mode searches (relative to nonlabile searches) of AI-ETD ADP-Ribosylation data, showing improved performance for combined searches with increasing laser power. B) ADP-ribosylated PSMs from labile, nonlabile, and combined searches of HCD-fragmented HeLa lysate. C) ADP-ribosylated PSMs from mouse liver tissue. D) Comparison of labile searches with −42 fragment remainder ion from HeLa lysate and mouse liver tissue, showing improved performance in mouse tissue but not HeLa. Pie charts show majority of mouse tissue ADP-ribosylation sites are at Arg residues (blue) vs majority unlocalized (red) in HeLa, likely corresponding to Ser residue sites.

## Conclusions

Labile mode search in MSFragger provides a powerful and flexible set of tools to identify spectra of peptides bearing labile modifications. While the degree of improvement in labile search compared to traditional, nonlabile search scales with the activation energy used to fragment the peptide and the lability of the modification, the combined-mode search option improves the spectrum annotation rate in nearly all cases by searching for peptides both with and without modification fragmentation. With user-specified options for diagnostic ions, peptide remainder masses, and fragment remainder masses, a wide variety of labile modifications can be easily searched with MSFragger. The labile mode search can leverage the fragment ions discovered by our recently described fragment ion discovery module in PTM-Shepherd(23), allowing labile mode to be used without requiring manual annotation of fragmentation pathways for new modifications. Localization of labile modifications has long been a significant obstacle to PTM analyses, and remains so in cases where PTMs dissociate without forming any fragment remainder masses. However, in many cases we can now localize modifications using fragment remainder ions, as demonstrated for RNA photo-crosslinking and Arg-linked ADP-ribosylation here. PTMProphet localization is also available in FragPipe and has the ability to localize using neutral loss masses, allowing for confirmatory localization of labile modifications following MSFragger search. Labile search mode is implemented in MSFragger version 3.5 and FragPipe version 18.0, along with visualization support in the integrated FP-PDV visualization software. Included with FragPipe are pre-built workflow for labile searches of phosphorylation and ADP-ribosylation, which can also easily be modified to allow user-friendly implementation of labile searches for any modification of interest.

**Table 1.** Phosphopeptides and proteins from CPTAC search. Combined and labile mode searches annotated ^~^4000 more phosphopeptides and 300 more phosphoproteins from the same raw data in the CPTAC TMT phosphoproteomics dataset.

| Search Type | Variable<br>Mods | Offsets | Phospho<br>PSMs | Phospho<br>peptides | Phospho<br>proteins |
| --- | --- | --- | --- | --- | --- |
| Nonlabile | 3 | 0 | 2,047,690 | 59,516 | 8,894 |
| Combined | 2 | 1 | 2,198,545 | 63,909 | 9,172 |
| Combined | 1 | 2 | 2,232,536 | 64,702 | 9,178 |
| Labile | 0 | 3 | 2,278,761 | 64,073 | 9,220 |

## Abbreviations

PTM: post-translational modification
MS: mass spectrometry
FDR: false discovery rate
PSM: peptide-spectrum match
HCD: higher-energy C-trap dissociation
AI-ETD: activation ion-electron transfer dissociation
NCE: normalized collision energy
RNA-XL: RNA-protein crosslink
4SU: 4-thiouridine
CCRCC: clear cell renal carcinoma
CPTAC: Clinical Proteomics Tumor Analysis Consortium
LC-MS/MS: liquid chromatography-tandem mass spectrometry

## Data Availability

Raw data reanalyzed here can be found in the public repositories listed in the Experimental Methods section (see “Datasets”). Processed tables containing results (PSMs, peptides, and proteins found) for all searches described here have been deposited at https://doi.org/10.5281/zenodo.7186868. FragPipe workflow files containing all parameters used for each search are also provided in this repository. A guide explaining the format of the results tables can be found at https://fragpipe.nesvilab.org/docs/tutorial_fragpipe_outputs.html.

## Acknowledgements

This work was funded in part by NIH grants R01GM094231, 5R01GM135504, and U24CA271037.

